# Polysaccharides from the fungus *Inonotus obliquus* activate macrophages into a tumoricidal phenotype via interaction with TLR2, TLR4 and Dectin-1a

**DOI:** 10.1101/2020.11.11.378356

**Authors:** CW Wold, PF Christopoulos, MA Arias, DE Dzovor, I Øynebråten, A Corthay, KT Inngjerdingen

**Author notes:** These authors contributed equally (A Corthay, KT Inngjerdingen).

## Abstract

**Background:** Tumor-associated macrophages (TAMs) may both promote and suppress tumor development. Therefore, molecules that are able to activate and repolarize these cells into a tumoricidal phenotype could be of great interest as a new strategy for cancer immunotherapy. Fungal β-glucans have been suggested as a promising way of activating TAMs, but most of the research has been carried out on particulate β-glucans of large sizes (> 500 kDa), which potentially have different biological properties than smaller, water-soluble molecules with similar structures.

**Methods:** Bone marrow-derived mouse macrophages were treated with 6 different purified polysaccharides isolated from the medicinal fungus *Inonotus obliquus*. Nitric oxide concentration was quantified using the Griess assay and by qPCR of iNOS mRNA. IL-6 and TNF-α concentrations were quantified using Luminex ELISA technology (using human monocyte-derived macrophages and mouse bone-marrow derived macrophages). Growth inhibition of cancer cells was measured using radiolabeled thymidine. Receptor interaction was determined using HEK-Blue™ reporter cell lines and TLR4 KO macrophages.

**Main findings:** The acidic, water-soluble polysaccharides AcF1, AcF2 and AcF3 induced nitric oxide (NO) production by mouse macrophages when combined with IFN-γ, leading to a strong subsequent tumoricidal activity by the macrophages. Tumoricidal activity of AcF1 and AcF3 was fully retained in TLR4 knockout macrophages, demonstrating that the macrophage activation was not dependent on TLR4. Further, AcF3 induced high levels of the pro-inflammatory cytokines IL-6 and TNF-α in human and mouse macrophages, independent of co-activation with IFN-γ. The polysaccharides were shown to bind TLR2, TLR4 and Dectin-1a to varying degrees, and these receptors were likely to be responsible for the macrophage activation. The acidic polysaccharides AcF1, AcF2 and AcF3 strongly activated TLR2, while AcF3 and AcF1 activated TLR4. The acidic polysaccharides had low affinity to Dectin-1a compared to the polysaccharides IWN, EWN and A1, which suggests that this receptor is not the main receptor for the pro-inflammatory activity observed.

**Conclusion:** For the first time, this study demonstrates that *I. obliquus* polysaccharides are able to bind multiple pattern recognition receptors to activate macrophages into a pro-inflammatory anti-tumor phenotype. The induction of tumoricidal activity in the macrophages as well as the interaction with TLR2, TLR4 and Dectin-1a suggest that the *I. obliquus* polysaccharides may have unique ways of interacting with macrophages, which could open up for new treatment options in cancer immunotherapy.

## Introduction

Cancer is a leading cause of mortality worldwide. The ability of our immune system to combat this complex and heterogeneous disease is well documented (1). Further, there is emerging evidence that certain immune cells such as macrophages can display both anti- and pro-tumorigenic phenotypes, depending on the microenvironment (2–4). For example, tumor-associated macrophages (TAMs) can act pro-tumorigenic by promoting angiogenesis to increase tumor growth and by dampening the general immune response against the tumor. In such cases, there is often a correlation between high TAM density and poor survival of cancer patients (5, 6). As such, macrophages are attractive targets for novel anti-tumor immunotherapy drugs, as exemplified by the recent increase in investments into this research field by the pharmaceutical industry (7). Polarization of macrophages into an anti-tumor “M1” phenotype by stimulation with toll-like receptor (TLR) ligands and interferon-γ (IFN-γ) causes upregulation of pro-inflammatory mediators such as nitric oxide (NO) and cytokines like tumor necrosis factor alpha (TNF-α) (8). This activation could induce an anti-tumor response by several distinct mechanisms, such as local tissue damage by NO, increased uptake of tumor neoantigens by the macrophages with subsequent presentation of the antigens to CD4^+^ T cells and CD8^+^ T cells, as well as activation of NK cells (3, 6, 9).

Fungal polysaccharides, especially β-glucans, have recently emerged as promising candidates for macrophage polarization into the M1 phenotype, due to their non-toxic nature and their potential to bind pattern recognition receptors (PRRs) such as Dectin-1 (10–12). Fungal β-glucans are large polymers consisting of glucose in a β-anomeric configuration. The consensus is that particulate, large-sized β-glucans with (1→3) and/or (1→6)-linkages between the monomers are able to elicit strong immune responses by binding to the C-type lectin receptor Dectin-1 found on macrophages and dendritic cells (13). However, there is much conflicting evidence in the literature regarding fungal β-glucans, often because alleged β-glucans are not actually pure β-glucans but rather a mix of various polysaccharides and proteins. For example, when the “β-glucan” zymosan was treated with various solvents to further purify the β-glucan part, TLR2 binding capability was lost while affinity for Dectin-1 was fully retained (14). The TLR2 binding capabilities of zymosan was attributed to other fungal compounds, and it now appears evident that smaller-sized, water-soluble fungal polysaccharides are able to bind other PRRs apart from Dectin-1, including TLR2 (15).

*Inonotus obliquus*, a white-rot fungus found on birch trees in the northern hemisphere, is a promising source of bioactive polysaccharides, and several authors have described immunomodulating polysaccharides from wild and cultivated sources of this fungus (16, 17). However, systematic characterization of the immunological properties of *I. obliquus* polysaccharides is lacking. Further, the receptors responsible for the activity have not yet been established. It has been suggested by some authors that TLR2 was involved in activation by a crude polysaccharide extract from *I. obliquus*, but the extract was not chemically characterized (18). We have previously isolated polysaccharides from the water- and alkali extracts from *I. obliquus* (19). We found that most of the polysaccharides had a main structural motif consisting of (1→3/1→6)-β-glucose, a typical| immunologically promising motif seen in fungal polysaccharides. Further, the water-soluble polysaccharides were complex and highly branched, and contained several types of monosaccharides in addition to glucose, such as (1→6)-α-galactose, (1→4)-α-galacturonic acid, (1→3)-α-mannose and (1→4)-β-xylose. The polysaccharides tested negative for lipopolysaccharide (LPS) contamination by GC-MS analysis (detection limit corresponded to 1.4 ng/mL LPS in a 100 μg/mL polysaccharide sample used for bioassays). The polysaccharides were screened using the Griess assay to measure NO production in the two cell lines J774A.1 and D2SC/1, and based on preliminary results, six polysaccharides were deemed promising for further immunological testing, and these form the basis of the studies presented herein. Here, we have performed an extensive immunological characterization of the isolated and purified *I. obliquus* polysaccharides, including their tumoricidal potential and which immune receptors they activate, with the goal to increase knowledge of the immunotherapeutic potential of this popular “edible” fungus.

## Materials & methods

### Isolation and characterization of polysaccharides from *I. obliquus*

Sclerotia (dense fungal conks) of *I. obliquus* were harvested from a birch tree in Oslo, Norway, and polysaccharides were isolated from the fungal material and characterized as previously described (19). Briefly, dried material from the interior part of *I. obliquus* (IOI) or the exterior part of *I. obliquus* (IOE) was boiled in water or 1 M NaOH, before the extracts were precipitated using 70 % EtOH and dialyzed (cut-off 3.5 kDa). The extracts were then fractionated using column chromatography. The water-soluble extracts (W) were first applied to an ion-exchange column using a NaCl gradient to create neutral (N) as well as acidic (Ac) fractions. Two neutral, water-soluble extracts were isolated, one from the interior part (IOI-WN) and one from the exterior part (IOE-WN). The acidic fraction IOI-WAc was separated further using a size-exclusion column to yield IOI-WAcF1, IOI-WAcF2 and IOI-WAcF3. The alkaline extract was applied to a size-exclusion column to yield IOI-A1. The fractions were characterized using several different methods, including gas chromatography (monosaccharide composition), gas chromatography-mass spectrometry (GC-MS), for linkage analysis), 2D NMR spectroscopy (anomeric configuration), size exclusion chromatography – multiple-angle laser light scattering (SEC-MALLS for molecular weights) and Smith degradation (general relationship between parts of the polymer).

### Mice

C57BL/6NRj mice were purchased from Janvier Labs (Le Genest-Saint-Isle, France) and bred at the Department of Comparative Medicine, Oslo University Hospital, Rikshospitalet (Oslo, Norway) in specific pathogen free (SPF) conditions. Bone marrow cells from C57BL/6NRj mice deficient in TLR4 (Tlr4^-/-^) were kindly provided by Maykel Arias, University of Zaragoza.

### Isolation and differentiation of bone marrow-derived macrophages

Conditioned medium (CM) containing macrophage colony-stimulating factor (M-CSF) was generated by culturing L929 cells in RPMI 1640 medium with L-Glutamine (Thermo Fisher Scientific) containing 10 % fetal bovine serum (FBS, Biochrom GmbH) and 1 % penicillin-streptomycin (P/S, Sigma-Aldrich). After 10 days, CM was collected, centrifuged, filtered and stored at −20°C, and was used for the differentiation and maintenance of bone marrow-derived macrophages (BMDMs). For isolation of the macrophages, femurs and tibiae of the hind legs from 8- to 12-week-old C57BL/6NRj mice and C57BL/10ScN TLR4^(-/-)^ knockout (KO) mice were harvested and flushed with RPMI 1640 medium containing 10 % FBS under sterile conditions. The bone marrow was passed through a cell strainer with 70 μm pores (Sigma-Aldrich) and cultured in non-tissue culture treated dishes (10 cm, VWR) in RPMI medium with L-Glutamine containing 10 % FBS, 1 % P/S and 30 % L929-derived CM. The cells were cultured for 5 days, after which non-adherent cells were washed off using phosphate buffered saline (PBS, with MgCl_2_ and CaCl_2_, Sigma-Aldrich) and the adherent macrophages were cultured for 2 more days. Macrophages were then harvested by incubation (20 min at 4°C) with cold PBS (without CaCl_2_ and MgCl_2_, Sigma-Aldrich). Macrophages were then flushed off the plate, collected, counted and kept frozen in aliquots at −150°C in FBS with 10 % dimethyl sulfoxide (DMSO, VWR) for future experiments. The purity of the cells was 99 % as analyzed by flow cytometry using the macrophage markers CD11b (M1/70, BioLegend) and F4/80 (BM8, BioLegend), and the cells were then referred to as wild-type (WT) or TLR4 KO bone marrow-derived macrophages (BMDMs).

### Isolation and differentiation of human monocyte-derived macrophages

Buffy coats were obtained from the Blood bank at Oslo University Hospital and approved for use by the Norwegian Regional Committee for Medical and Health Research Ethics, REK no. 2019/113. The buffy coats were mixed with an equal volume of PBS containing 2 % FBS, before gently added on to Lymphoprep™ in 50 mL tubes in volumes recommended by the provider (Progen, #1114545; Heidelberg, DE). The tubes were centrifuged at 800 g for 20 min at room temperature (RT), before the middle, buffy layer containing peripheral blood mononuclear cells was collected and washed twice in PBS by centrifugation (400 g, 7 min, RT). Next, the cells were filtered through a 30 μm filter to remove cell clumps and debris, before the monocytes were positively selected by magnetic-activated cell sorting (MACS) technology using CD14 MicroBeads (Miltenyi) according to the manufacturer’s instructions. Briefly, magnetic beads conjugated to an anti-human CD14 antibody (CD14 MicroBeads) were added to the peripheral blood mononuclear cells and incubated for 15 min at 4 °C. Unbound, excess beads were washed away, and the cell suspension was applied to a column placed in a MACS magnetic separator. Unlabeled cells were washed out before the column was removed from the magnet field and the magnetically labeled cells were flushed out with a plunger. Staining with APC/Cy7-conjugated anti-human CD14 antibody (clone HCD14, BioLegend) followed by flow cytometry, showed that > 95% of the positively selected cells were monocytes. APC/Cy7-conjugated mouse IgG1k (clone MOPC-21, BioLegend) was used as an isotype-matched control antibody.

The positively selected CD14^+^ monocytes were differentiated into macrophages by cultivation for 6 days in medium with granulocyte-macrophage colony-stimulating factor (GM-CSF, Peprotech). More specifically, 3 x 10^6^ cells were seeded out in 10 mL RPMI 1640 containing 10 % FBS, 1 % P/S, and 50 ng/mL GM-CSF per 10 cm non-tissue culture treated dish (VWR). At day 3, half of the medium was replenished with fresh medium containing 50 ng/mL GM-CSF. To harvest the macrophages on day 6, the dishes were put on ice for 30 min, while polypropylene falcon tubes (50 mL, SARSTEDT) were pre-coated with FBS by swirling the tubes carefully before put on ice. Cell culture medium was harvested from the dishes and collected in the pre-coated falcon tubes. Next, ice-cold PBS with 2.5 mM EDTA was added to the dishes for 10 min before the macrophages were detached and collected by pipetting up and down several times. A cell scraper was used to detach the remaining macrophages.

### Pattern recognition receptor agonists and cytokines

The following PRR agonists were used as controls in various experiments: Pam_3_CysSerLys4 (Pam_3_CSK_4_, TLR1/TLR2 agonist, InvivoGen); Lipopolysaccharide (LPS) from *Escherichia coli* and *Salmonella minnesota* (both ultrapure TLR4 agonists, InvivoGen); Lipoteichoic acid (LTA) from *Staphylococcus aureus* (TLR2/TLR6 agonist, Sigma-Aldrich); Cl_2_64 (TLR7 agonist, InvivoGen), zymosan from *Saccharomyces cerevisiae* (zymosan crude/ZymC, TLR2 and Dectin-1 agonist, Sigma-Aldrich), zymosan depleted from *S. cerevisiae* (zymosan purified/ZymP, InvivoGen), laminarin from *Laminaria digitata* (Dectin-1 ligand, Sigma-Aldrich). The PRR agonists were used alone or in combination with 20 ng/mL mouse recombinant IFN-γ (Peprotech).

### Quantification of nitric oxide

NO production by activated macrophages was measured using the Griess reagent system as previously described with some modifications (19). Cells were seeded out in a flat bottom 96-well plate (Costar) at cell density 6 × 10^4^ cells to a final volume of 200 μL/well. Cell medium was RPMI 1640 with L-Glutamine containing 10 % FBS and 10 % CM. After 24 h treatment with *I. obliquus* polysaccharides or other activation factors, cell media (100 μL) were collected and centrifuged (400 *g*, 2 min), and 50 μL supernatant was then mixed with equal parts Griess reagents A (dH_2_O with 1 % sulphanilamide [Sigma-Aldrich] and 5 % phosphoric acid [Sigma-Aldrich]). The mixture was incubated in the dark for 10 min at room temperature (RT), before 50 μL of Griess reagent B (0.1 % *N*-(1-napthyl) ethylenediamine [Sigma-Aldrich] in dH_2_O) was added in order to convert NO into nitrite (NO_2_^-^), which was quantified colorimetrically at A_540_ using NaNO_2_ (1.56 – 100 μM) as a standard curve. Samples were set up in duplicates or triplicates depending on the experimental setup. For experiments using the LPS inhibitor polymyxin B (PMB, Polymyxin B sulfate salt, Sigma-Aldrich), the samples were mixed with PMB and incubated for 30 min at RT before being added to the cultivated cells. The experiments were carried out at least three times.

### Tumor Cell Growth Inhibition Assay

Wild-type (WT) and TLR4 knockout (KO) BMDMs were thawed and cultured for 3 days in non-Stissue culture treated dishes (VWR) in RPMI 1640 medium with L-Glutamine containing 10 % FBS and 10 % CM (= complete medium). The BMDMs were harvested by scraping, incubated for 2 h at 37°C with mitomycin C (10 mg/mL, Sigma-Aldrich) to inhibit proliferation, and then washed twice with PBS. Next, the BMDMs were resuspended in the same type of medium and seeded out in triplicates in flat bottom 96-well plates (Costar) at three densities: 6 × 10^4^, 3 × 10^4^, and 3 × 10^3^ cells/well in a final volume of 200 μL/well. After 20 h, half of the medium was replaced with complete medium containing *I. obliquus* polysaccharides with or without IFN-γ, and incubated for 24 h. Next, half of the cell supernatants (100 μL) were removed and used for quantification of NO_2_-. Lewis lung carcinoma (LLC) cells (CLS Cell Line Service, 3 × 10^3^ cells/well) were then added to the macrophages, resulting in varying ratios of effector to target cells: 20:1, 10:1 and 1:1. After 20 h of co-culture, ^3^H-thymidine (10 μL, 0.2 μCi/well, Hartmann Analytic) was added and the cells were harvested 24 h later after a freeze- and thaw cycle. The amount of radiolabeled DNA was measured on a 1450 MicroBeta Trilux Microplate Scintillation counter (Perkin Elmer). The experiments were carried out at least three times.

### Determination of iNOS mRNA Levels by Real-Time Quantitative PCR

BMDMs were seeded in 12-well plates (Sigma-Aldrich) at a density of 6 × 10^5^ cells/well in 1 mL RPMI medium with L-Glutamine supplemented with 10 % FBS and 10 % CM. The cells were incubated for 2h, before 0.5 mL of the medium was removed and replaced with 0.5 mL medium containing *I. obliquus* polysaccharides with or without IFN-γ (20 ng/mL). After 24 h, cell culture media were removed and total RNA was extracted from the cells by using 300 μL/well of TRI Reagent (Merck) and Direct-zol RNA minipreps (Zymo Research) according to manufacturer’s instructions. Next, mRNA concentrations were measured using Nanodrop One/One (Thermo Fisher), and 250 ng RNA of each sample was reverse transcribed to cDNA using the Primescript RT kit (Takara Bio) according to the manufacturer’s instructions. Real-time quantitative PCR (qPCR) was performed with 50 ng of the obtained cDNA, using a Kapa SYBR fast qPCR kit (Kapa Biosystems) and 0.2 μM of mRNA specific primers for the mouse gene *Nos2* which encodes iNOS (forward primer: TTCACCCAGTTGTGCATC GACCTA, reverse primer: TCCATGGTCACCTCCAACACA AGA) and with primers for 18s rRNA (forward primer: CGCTTCCTTACCTGGTTGAT, reverse primer: GAGCGACCAAAGGAACCATA) as the endogenous control, at temperature cycling conditions: 95°C for 3min, then 95°C for 3 s and 60°C for 30 s for 40 cycles. All samples were run in duplicates and the final values were averaged. Following melting curve analysis, the relative differences in iNOS mRNA levels were expressed using the – ΔCq values (Cq 18s rRNA - Cq iNOS), where a more negative value means lower relative expression of iNOS mRNA compared to the housekeeping gene 18S rRNA. One unit increase in the negative ΔCq value corresponded to a doubling of iNOS mRNA. The experiment was carried out three times.

### Quantification of pro-inflammatory cytokines using Luminex technology

Mouse BMDMs, 2.5 x 10^5^ cells in 0.5 mL medium per well in 24-well plates (Costar) were cultured in RPMI 1640 supplemented with 10 % FBS (Biochrom) and 10 % CM. Human monocyte-derived macrophages, 1 x 10^5^ cells in 0.3 mL medium per well in 48-well plates (Costar) were cultured in RPMI 1640 supplemented with 10 % FBS (Biowest) and stimulated for 24 h with polysaccharides with or without IFN-γ. Cell culture media were collected and centrifuged (1000g for 15 min at 4°C). Next, the supernatants were moved to new Eppendorf tubes and centrifuged again (1000g for 15 min at 4°C) to remove cells and debris before storage at −80°C until analysis. The concentrations of mouse and human IL-6 and TNF-α were determined by a multiplex Bio-Plex assay (Bio-Rad) according to the manufacturer’s instructions. Samples were analyzed in duplicates, using a Bio-Plex MAGPIX Multiplex Reader and Bio-Plex Manager 6.1 software (Bio-Rad Laboratories). The experiment was carried out three times.

### Macrophage proliferation assay

BMDMs were seeded out in triplicates in a flat bottom 96-well plate (Costar) at cell density 6 × 10^4^ cells to a final volume of 200 μL/well. The cells were left to attach, and after 2 h, the medium was replaced with medium containing *I. obliquus* polysaccharides with or without IFN-γ, and incubated for 24 h. Then ^3^H-thymidine (10 μL, 0.2 μCi/well, Hartmann Analytic) was added and after another 24 h the cells were harvested after a freeze- and thaw cycle. The amount of radiolabeled DNA was measured on a 1450 MicroBeta Trilux Microplate Scintillation counter (Perkin Elmer). The experiment was carried out three times.

### Reporter cell lines

HEK-Blue™ reporter cell lines (Invivogen) transfected with human TLR2, human Dectin1a, human TLR4/CD14/MD2 or non-transfected (null-1) were cultured and maintained using DMEM GlutaMAX™ containing 10 % FBS (Sigma-Aldrich), 1 % P/S, Normocin (100 μg/mL) and HEK-Blue™ selection antibiotics. Experiments were carried out according to the manufacturer’s instructions. Briefly, *I. obliquus* polysaccharides (20 μL) at various concentrations were added to wells in 96-well plates (Costar). Then, cells were gently washed with warm PBS before suspended in HEK-Blue™ SEAP detection medium. Finally, the cells were seeded out in a density of 5 x 10^4^ cells/well to the wells containing samples and detection medium. After 16 h incubation (37 °C, 5 % CO_2_), secreted alkaline phosphatase (SEAP) was detected colorimetrically at A_635_. The experiments were carried out at least three times.

### Statistical analysis

Statistical analysis was conducted by using the GraphPad Prism 8 software (GraphPad). Values from control samples were used as a reference to each individual value, unless otherwise stated. The data were analyzed using one-way ANOVA test, followed by Dunnett’s test for comparison of multiple samples, or using two-way ANOVA followed by Sidak’s test for comparison of multiple samples. The values were compared either across the data set or individually against the controls depending on the experiment (stated specifically below each experiment figure).

## Results

### Polysaccharides from *I. obliquus* synergize with IFN-γ to induce nitric oxide production by macrophages in a dose-dependent manner

The main structural and chemical characteristics of the *I. obliquus* polysaccharides used in this study are summarized in Table 1, and simplified structural models of the polysaccharides based on GC and GC-MS analysis and NMR spectroscopy are shown in Figure 1. Although the drawings in Figure 1 represent probable structural compositions, they do not necessarily reflect the structural complexity or the accurate relationship between the monomers. For detailed characteristics of the polysaccharides, see Wold et al (2018). Based on their structural motifs and molecular weight, the polysaccharides could be allocated to three main structural types, defined as neutral and water-soluble (IWN and EWN), acidic and water-soluble (AcF1, AcF2 and AcF3) or neutral and particulate (A1).

**Table 1:** Names (abbreviations in parenthesis) and chemical characteristics of the polysaccharides from *I. obliquus* used in this study.

| **Polysaccharide fraction**<br>**(incl. abbreviations)** | **Water-**<br>**solubility** | **Mol. weight**<br>**(kDa)** | **pH**<br>**sensitivity** | **Main monomeric units** |
| --- | --- | --- | --- | --- |
| **IOI-WN**<br>**(IWN)** | High | 60 | Neutral | (1→3/1→6)-β-glucose<br>(1→6)-α-galactose |
| **IOE-WN**<br>**(EWN)** | High | 73 | Neutral | (1→3/1→6)-β-glucose<br>(1→6)-α-galactose |
| **IOI-WAcF1**<br>**(AcF1)** | High | 28 | Acidic | (1→3/1→6)-β-glucose<br>(1→6)-α-galactose<br>(1→4)-α-galacturonic acid |
| **IOI-WAcF2**<br>**(AcF2)** | High | 14 | Acidic | (1→3/1→6)-β-glucose<br>(1→6)-α-galactose<br>(1→4)-α-galacturonic acid<br>(1→4)-β-xylose |
| **IOI-WAcF3**<br>**(AcF3)** | High | 10 | Acidic | (1→3/1→6)-β-glucose<br>(1→6)-α-galactose<br>(1→4)-α-galacturonic acid<br>(1→4)-β-xylose |
| **IOI-A1**<br>**(A1)** | No (particulate) | >450 | Alkaline-extracted | (1→3/1→6)-β-glucose<br>(1→4)-β-xylose |

Abbreviations/names: IOI = *I. obliquus* Interior part; IOE = *I. obliquus* Exterior part; WN = Water-extracted, Neutral; WAc = Water-extracted, Acidic; F1-F3 = Fraction 1-3; A1 = Alkaline-extracted polysaccharide 1

**Figure 1:**
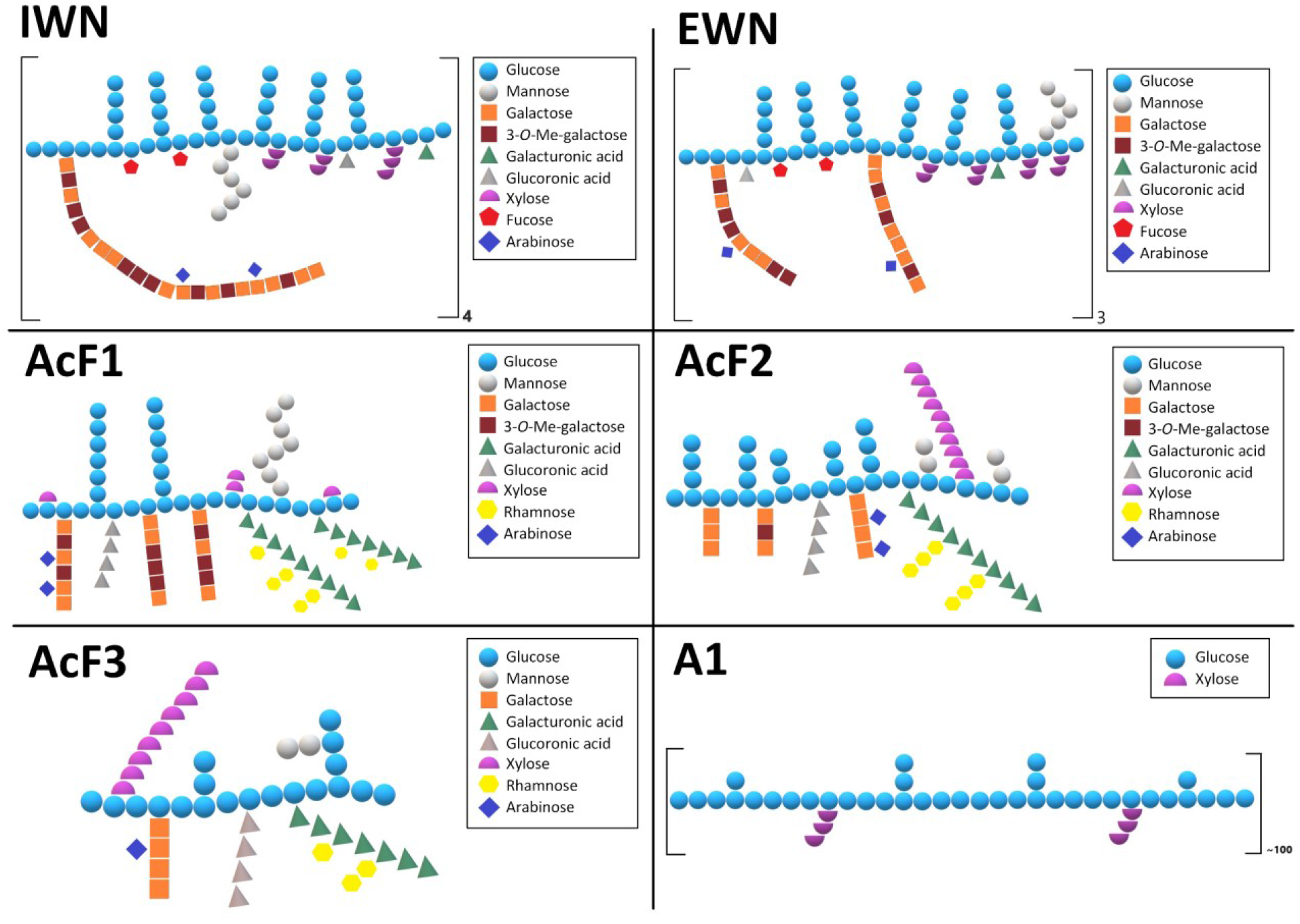
Tentative structures of the polysaccharides isolated from *I. obliquus*. The structures presented here are based on data from GC, GC-MS and NMR spectroscopic data. The drawings are theoretical possibilities and do not necessarily reflect the three-dimensional structure of the polysaccharides or the accuracy of the connection between the different monomers. IWN and EWN are neutral polymers, with β-glucose and α-galactose being the main monomeric units. AcF1, AcF2 and AcF3 are acidic polysaccharides with β-glucose as their main monomeric unit, with varying amounts of α-galacturonic acid, α-galactose and β-xylose. A1 is an alkaline polysaccharide with much higher molecular weight than the rest, and consists almost exclusively of β-glucose, in addition to minor amounts of β-xylose.

To investigate if the polysaccharides were able to activate macrophages, bone marrow-derived mouse macrophages (BMDMs) were generated from C57BL/6NRj mice. The capability of the polysaccharides to activate these cells was first measured by their ability to induce nitric oxide (NO) production by the macrophages, seen indirectly by levels of nitrite in the cell supernatant using the Griess assay (Figure 2A). NO production is an important marker of a pro-inflammatory M1-like macrophage phenotype (6). BMDMs were incubated with polysaccharides alone or in combination with IFN-γ, which is known to synergize with Toll-like receptor (TLR) ligands to induce a pro-inflammatory antitumor macrophage phenotype (8). As seen in Figure 2A, the polysaccharides were capable of activating macrophages in a dose-dependent manner when used in combination with IFN-γ. Most of the polysaccharides required co-activation with IFN-γ to activate the macrophages; however, in the absence of IFN-γ, AcF1 statistically significantly increased the NO production and AcF3 clearly showed a trend toward increased NO production, and these two polysaccharides were more potent than the rest when used in combination with IFN-γ. Nitrate levels in the cell supernatant is an indirect way of observing NO activation, because it requires the conversion of NO to nitrite using Griess reagents A and B. Another way of observing NO production by the macrophages is to quantify inducible nitric oxide synthase (iNOS) mRNA, the gene responsible for producing NO from L-arginine in the cells (20). Figure 2B shows iNOS mRNA levels compared to 18S rRNA after the macrophages were activated with polysaccharides at 100 μg/mL with or without IFN-γ. The iNOS results correlate with nitrite levels seen in Figure 2A, thus confirming that the polysaccharides were able to induce NO production in mouse macrophages.

**Figure 2:**
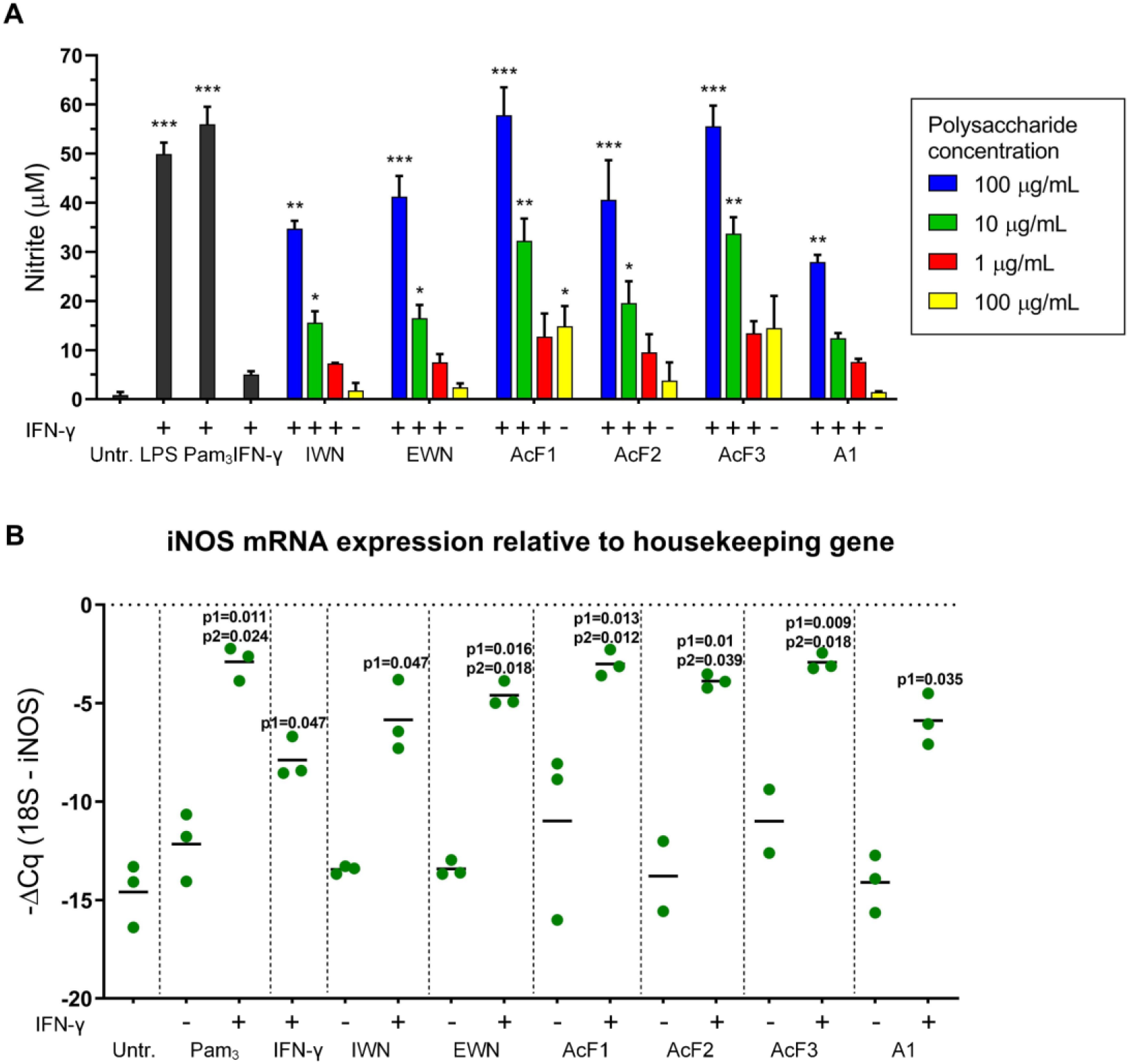
Polysaccharides synergize with IFN-γ to induce NO production in mouse macrophages. **A)** Bone marrow-derived macrophages (BMDMs) were incubated for 24 h with polysaccharides at different concentrations (1, 10 and 100 μg/mL) with IFN-γ (20 ng/mL), or at 100 μg/mL alone, before nitrite levels in the supernatant were quantified using the Griess assay. Lipopolysaccharide (LPS) from *E. coli* (100 ng/mL) and the TLR ligand Pam_3_-CSK_4_ (100 ng/mL) together with IFN-γ (20 ng/mL) were used as positive controls. Three independent experiments were performed, and the results are shown as means ± SD. Statistical significance was calculated using one-way ANOVA followed by Dunn’s multiple comparison test, with single comparisons against the untreated control *** = p < 0.001, ** = p < 0.01, * = p < 0.05. **B)** BMDMs were incubated for 24 h with polysaccharides (100 μg/mL) with or without IFN-γ (20 ng/mL), before cells were harvested, mRNA was isolated and iNOS mRNA and 18S rRNA were quantified using RT-qPCR. Pam_3_-CSK_4_ (100 ng/mL) and IFN-γ (20 ng/mL) alone or in combination were used as controls. Each dot in the graph represents the mean value of technical triplicates from one experiment. The black lines indicate means for two or three independent experiments. Statistical significance was calculated using one-way ANOVA followed by Dunn’s multiple comparison test, with single comparisons against either the untreated control (p1) or against the IFN-γ treated samples (p2).

### BMDMs stop proliferating upon activation by polysaccharides in combination with IFN-γ

Macrophages are capable of quickly reprogramming themselves into a pro-inflammatory phenotype upon recognition of pathogen-associated molecular patterns (PAMPs) such as LPS (21). This reprogramming is regarded as a committed step in macrophage polarization, because it requires the cells to exit their proliferative state and shift their metabolic resources into producing pro-inflammatory mediators to handle the incoming threat (22). Therefore, we wanted to explore if activation by *I. obliquus* polysaccharides could affected the macrophages’ proliferation capabilities, indicative of a pro-inflammatory phenotype. An experiment was conducted by treating BMDMs with radiolabelled thymidine after 24 h of treatment with *I. obliquus* polysaccharides to measure thymidine DNA incorporation in dividing cells. As seen in Figure 3, the proliferation of BMDMs was drastically reduced after incubation with polysaccharides in combination with IFN-γ. IWN, EWN, AcF2 or A1 alone did not result in any macrophage growth inhibition, but when combined with IFN-γ, the polysaccharides statistically significantly inhibited growth compared to the untreated cells. Interestingly, AcF1 and AcF3 inhibited proliferation to some extent on their own, similarly to LPS from *E. coli* (LPS-EK) and zymosan crude, but the effect was modest compared to growth inhibition measured after co-treatment with IFN-γ, which gave a strong inhibitory effect on proliferation. Thus, the results suggest that all the tested *I. obliquus* polysaccharides were able to induce extensive metabolic reprogramming in the macrophages by making them suppress proliferation, which is indicative of a pro-inflammatory phenotype.

**Figure 3:**
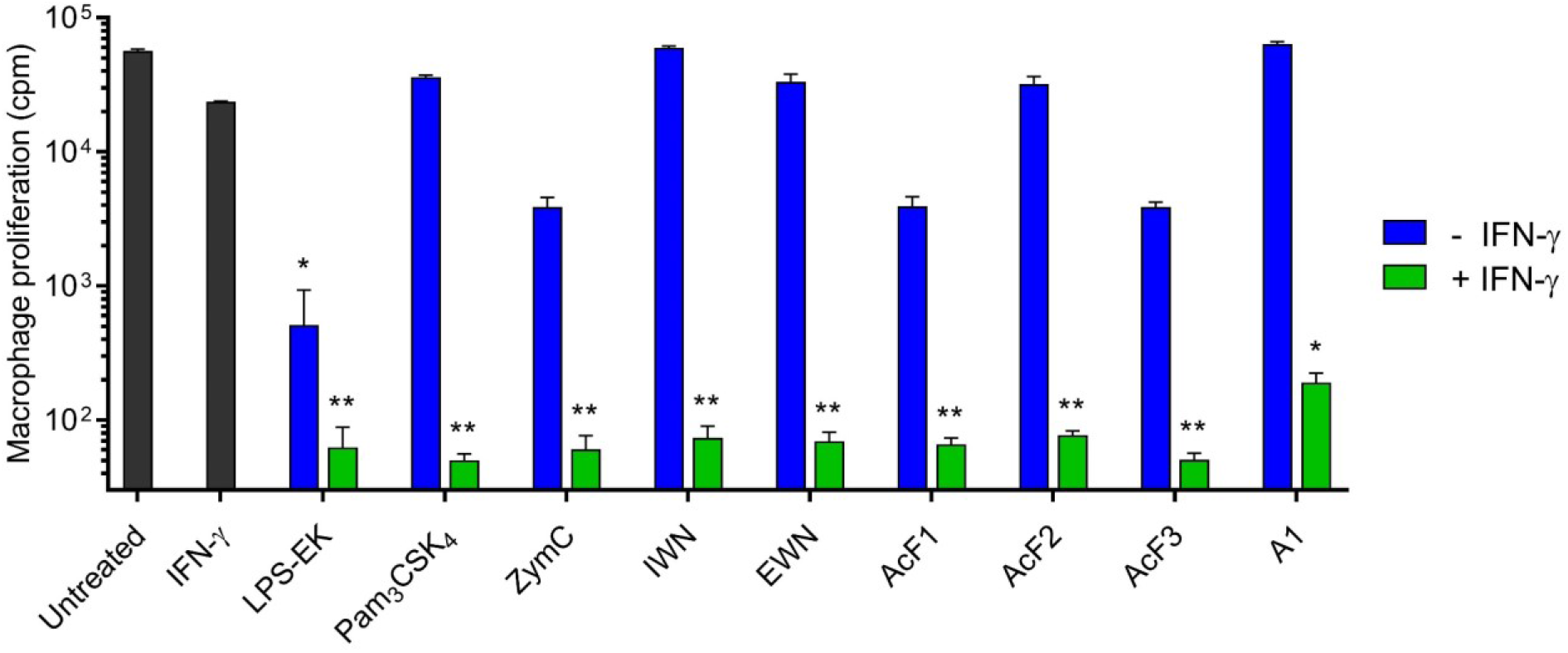
Polysaccharides from *I. obliquus* synergize with IFN-γ to inhibit macrophage proliferation. The cells were stimulated for 24 h before radiolabelled thymidine was added, and counted after another 24 h of incubation. Polysaccharides from *I. obliquus* were used at 100 μg/mL, LPS-EK (100 ng/mL), Pam_3_CSK_4_ (100 ng/mL), zymosan crude (ZymC, 100 μg/mL) and IFN-γ (20 ng/mL) were used as controls. Three independent experiments were performed, and average values ± SD are shown. Statistical significance was calculated using one-way ANOVA followed by Dunn’s comparison test, with single comparisons against the untreated control *** = p < 0.001, ** = p < 0.01, * = p < 0.05.

### Several *I. obliquus* polysaccharides synergize with IFN-γ to make macrophages inhibit cancer cell growth *in vitro*

To investigate the potential of *I. obliquus* polysaccharides to inhibit cancer cell growth, an *in vitro* growth inhibition assay was conducted by co-culturing macrophages with the Lewis lung carcinoma (LLC) cell line. As shown in Figure 4A, BMDMs were first treated with the DNA crosslinker mitomycin C to prevent their proliferation. Then, the macrophages were activated by polysaccharides and IFN-γ, before LLC cells were added to wells containing macrophages, in three different ratios between macrophages and cancer cells (20:1, 10:1 and 1:1). Ultimately, proliferation of cancer cells was measured after the addition of radiolabelled thymidine. Reduction of growth in this assay is an indirect measurement of macrophage-mediated cytostatic and/or cytotoxic effects on the cancer cells. Several of the polysaccharides were able to activate the macrophage as measured by the NO in the cell supernatants (Figure 4B), and this activation caused a strong inhibition of cancer cell proliferation (Figure 4C). All polysaccharides expect A1 gave a dose-dependent inhibitory effect, with the most prominent effect seen from AcF1 and AcF3 in combination with IFN-γ. The growth inhibition was almost equally effective at 10:1 as at 20:1, although there were some exception such as for AcF3 at 100 μg/mL where the 20:1 ratio was most effective. At the 1:1 ratio, the cancer cell proliferation was only marginally inhibited (data not shown). The results demonstrate the ability of the polysaccharides to activate macrophages into a tumoricidal phenotype. Further, NO concentration correlated strongly with the inhibitory effect, suggesting that NO is important for the cancer cell growth inhibition (8).

**Figure 4:**
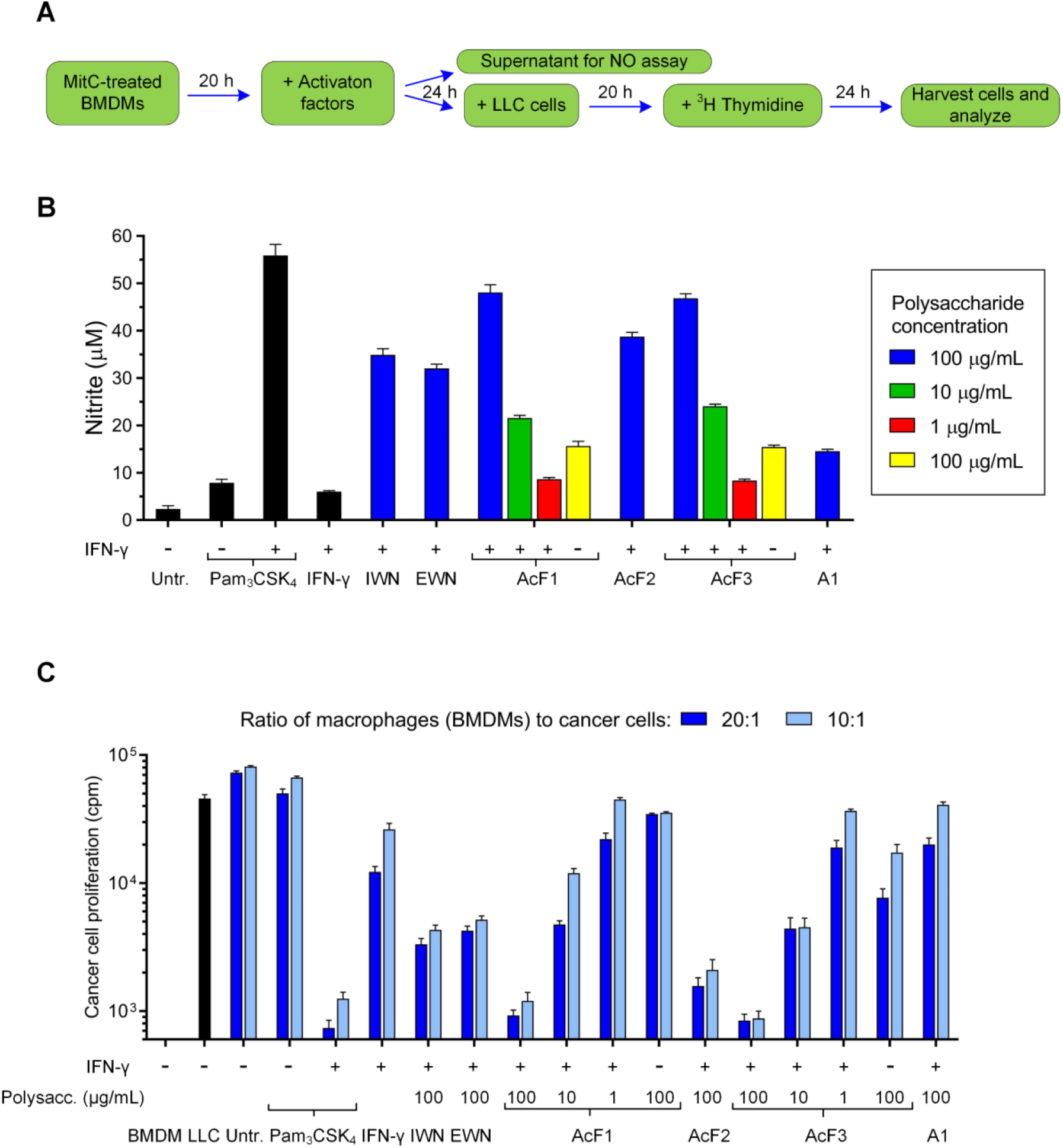
Polysaccharides from *I. obliquus* in combination with IFN-γ inhibit cancer cell growth by activating macrophages. **A**) The experimental setup: BMDMs (6 x 10^4^ cells/well or 3 x 10^4^ cells/well for 20:1 and 10:1 ratios, respectively) were incubated with mitomycin C for 2 h, then cultivated for 20 h before treatment with polysaccharides and/or IFN-γ for 24 h. Then, 100 μL cell medium was removed and analysed for NO before Lewis lung carcinoma (LLC) cells (3 x 10^3^ cells/well) were added to the well containing macrophages, giving two different ratios between macrophages and cancer cells (20:1 and 10:1). After 20 h, radiolabelled thymidine was added and the cells were incubated for another 24 h before inhibition of cancer cell growth was measured. **B)** NO concentration in the cell medium was measured using the Griess assay. The polysaccharide concentration was 100 μg/mL except for AcF1 and AcF3, which were tested in concentrations 1, 10 and 100 μg/mL. All samples were tested alone and in combination with IFN-γ (20 ng/mL). Pam_3_CSK_4_ (100 ng/mL) and IFN-γ (20 ng/mL) were used as controls. **C)** Cancer cell growth inhibition was measured using radiolabelled thymidine, expressed as radioactive counts per minute (cpm) values. The polysaccharide concentration was 100 μg/mL except for AcF1 and AcF3, which were tested in concentrations 1, 10 and 100 μg/mL. All samples were tested alone and in combination with IFN-γ (20 ng/mL). Pam_3_CSK_4_ (100 ng/mL) and IFN-γ (20 ng/mL) were used as controls. Untreated control (Untr.) consisted of co-cultured, non-activated BMDMs and LLC cells. Three independent experiments were performed, and a representative experiment is shown using average values ± SD from technical triplicates.

### The polysaccharides activate macrophages independently of LPS

LPS is a common contaminant when working with compounds of natural origin (23). Therefore, we wanted to rule out the possibility that our polysaccharide fractions were contaminated with LPS. LPS from *E. coli* can activate mouse macrophages at a concentration as low as 500 pg/mL (data not shown) and could thus be responsible for the observed bioactivity, even though LPS was not detected by GC-MS analysis of the polysaccharides (detection limit corresponded to 1.4 ng/mL for a 100 μg/mL sample) (19). We performed an experiment using the established LPS inhibitor polymyxin B (PMB) (24) which neutralizes LPS by binding to the Lipid A moiety, and thereby block potential LPS-mediated activation. *I. obliquus* polysaccharides and pattern recognition receptor (PRR) ligands were either kept untreated or were incubated with PMB before being added to BMDMs in combination with IFN-γ. Ultimately, NO concentration in cell media was measured using the Griess assay. As shown in Figure 5A, the PMB treatment did not influence the activity of AcF1, AcF2, AcF3 or A1 or the non-TLR4 PRR ligands used as controls. PMB treatment reduced the activity of IWN and EWN slightly, but most of the activity was retained in these samples as well. Two different types of LPS were used in the experiment: LPS from *S. minnesota* (LPS-SM) and LPS-EK (Fig. 5B). For both types of LPS, there was a complete inhibition of NO when LPS was pre-treated with PMB and used without co-treatment with IFN-γ. When LPS-SM and IFN-γ were used in combination, there was a ~50 % reduction in NO concentration at 100 ng/mL LPS-SM, ~80 % reduction at 10 ng/mL, and complete inhibition at 1 ng/mL. When LPS-EK and IFN-γ were used in combination, there was a ~75 % reduction in NO concentration when using 100 ng/mL LPS-EK, and at 10 ng/mL and 1 ng/mL the activity was completely inhibited. Because PMB was able to completely inhibit LPS at 1 ng/mL, and all the polysaccharides contained less than 1.4 ng/mL LPS at 100 μg/mL according to GC-MS analysis, the findings suggest that any potential LPS contamination in our samples should be completely blocked by PMB. Therefore, it can be concluded that there was no LPS present in the PMB-treated polysaccharide fractions AcF1, AcF2, AcF3 or A1. For IWN and EWN, a potential LPS contamination could not be ruled out, although the contamination would be minor as most of the activity of the fractions was retained after treatment with PMB.

**Figure 5:**
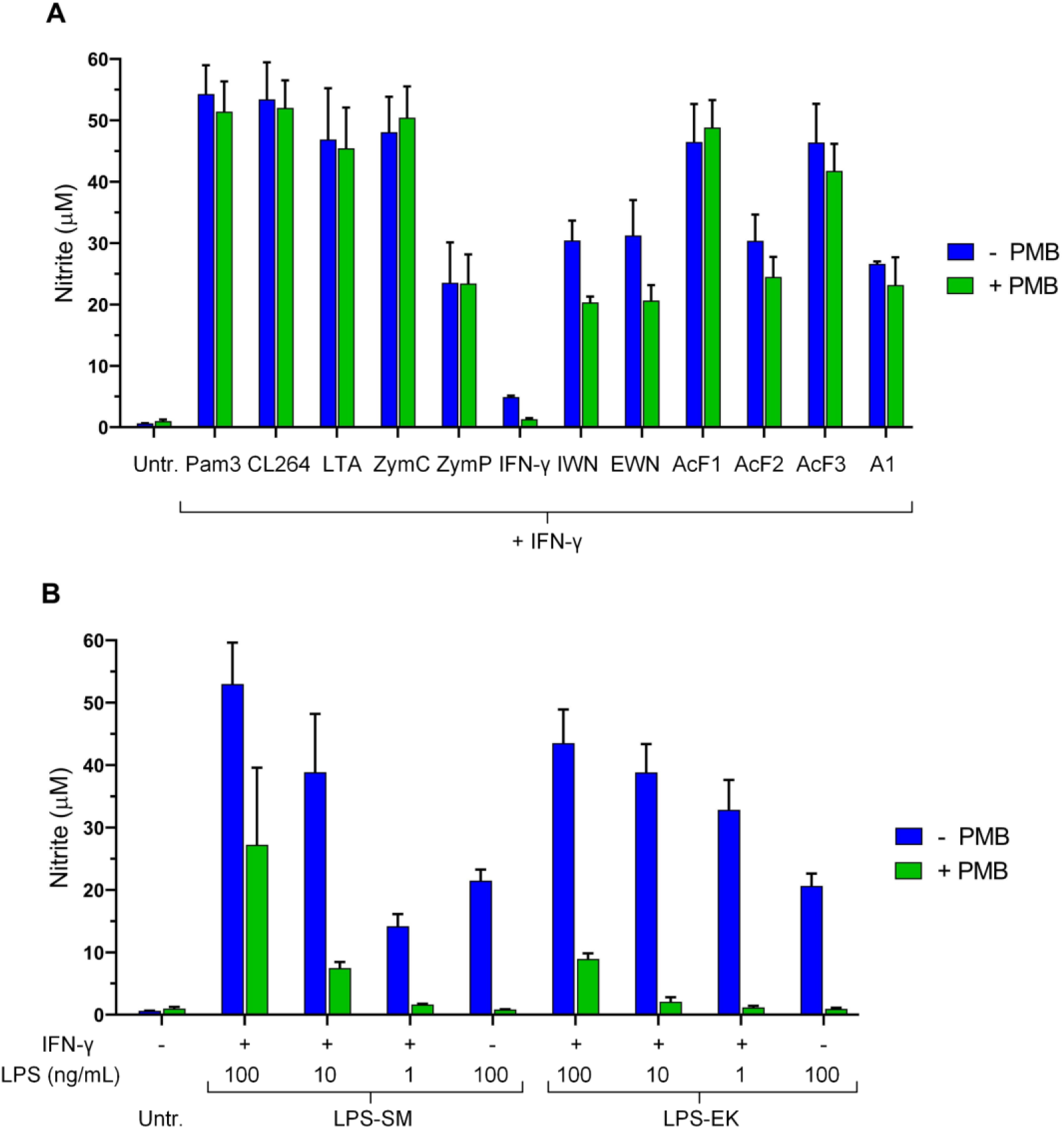
Polymyxin B (PMB) treatment did not inhibit *I. obliquus* polysaccharides from activating BMDMs, while LPS was strongly inhibited by the same treatment. PMB (10 μg/mL final concentration) was incubated with samples for 30 min, before the samples were added to BMDMs and incubated for 24 h, before NO concentration in the cell media was measured using the Griess assay. **A**) Polysaccharides from *I. obliquus* or various PRR ligands were able to activate BMDMs in combination with IFN-γ, even in the presence of PMB. Polysaccharides from *I. obliquus* were used at 100 μg/mL. Controls used: Pam_3_CSK_4_ (Pam_3_, 100 ng/mL, TLR1/2 agonist); Cl_2_64 (1 μg/mL, TLR7 agonist); LTA (100 μg/mL, TLR2 agonist); zymosan crude (ZymC, 100 μg/mL, TLR2/Dectin-1 agonist); zymosan purified (ZymP, 100 μg/mL, Dectin-1 agonist); IFN-γ (20 ng/mL). **B**) LPS-induced activation of BMDMs was inhibited by PMB treatment in a dose-dependent manner. LPS from *S. minnesota* (LPS-SM) and *E. coli* (LPS-EK) were used at 1, 10 and 100 ng/mL in combination with IFN-γ (20 ng/mL), or at 100 ng/mL alone. Three independent experiments were performed, and average values ± SD are shown. **A** and **B** are from the same experiments, the graphs are shown separately to highlight how PMB affects LPS in a dose-dependent manner.

### AcF1, AcF2 and AcF3 induce significant growth inhibition of cancer cells trough activation of TLR4 KO macrophages

Next, in order to confirm that the activation of the macrophages was mediated by the polysaccharides and not LPS contamination, BMDMs were generated from C57BL/6 TLR4^(-/-)^ (TLR4 knockout, KO) mice, as TLR4 is the receptor for LPS (25). The difference in activation between KO and wild type (WT) BMDMs was then compared after treatment with *I. obliquus* polysaccharides and IFN-γ. This was done by measuring NO concentration in cell supernatants by the Griess assay, as well as by using a growth inhibition assay with BMDMs co-cultured with LLC cells. As shown in Figure 6A, LPS-EK and LPS-SM were completely incapable of activating TLR4 KO macrophages to produce NO, whereas AcF1 and AcF3 retained their activity in these cells, with similar NO levels as in the WT cells. This demonstrates that AcF1 and AcF3 were able to activate macrophages through a different receptor than TLR4, and the potency of these polysaccharides in the KO cells was similar to the positive controls Pam_3_CSK_4_ and zymosan crude (ZymC). IWN, EWN and AcF2 had statistically significantly lower activity in the knockout cells. The NO concentration correlated mostly with growth inhibition (Figure 6B). Again, AcF1 and AcF3 were the most active polysaccharide fractions, with no significant differences between KO and WT cells. EWN had significantly reduced activity in the KO cells compared to the WT cells, while IWN and AcF2 had slightly reduced activity in the KO cells, although the difference was not statistically significant. AcF2 still gave a marked growth inhibition using both cell types. Interestingly, zymosan purified (ZymP) – a particulate β-glucan known as a specific Dectin-1 agonist – gave significantly increased inhibition of cancer cell growth when using TLR4 KO cells compared to WT cells, although the NO production was similar using the two cell types. The results presented in Figure 6B show a macrophage to cancer cell ratio of 10:1, which gave similar effects as the 20:1 ratio. At 1:1 the inhibitory effect was lost (data not shown). Because the main goal of this experiment was to investigate differences between WT and TLR4 KO cells, the statistical significances presented in Figure 6 are given as differences between WT and TLR4 KO macrophages within each sample. The results are in agreement with the results presented in Figure 5 using the LPS inhibitor PMB, and demonstrates the ability of the polysaccharides AcF1, AcF2 and AcF3 to activate macrophages into a tumoricidal phenotype independently of TLR4 and thereby of any putative LPS contamination.

**Figure 6:**
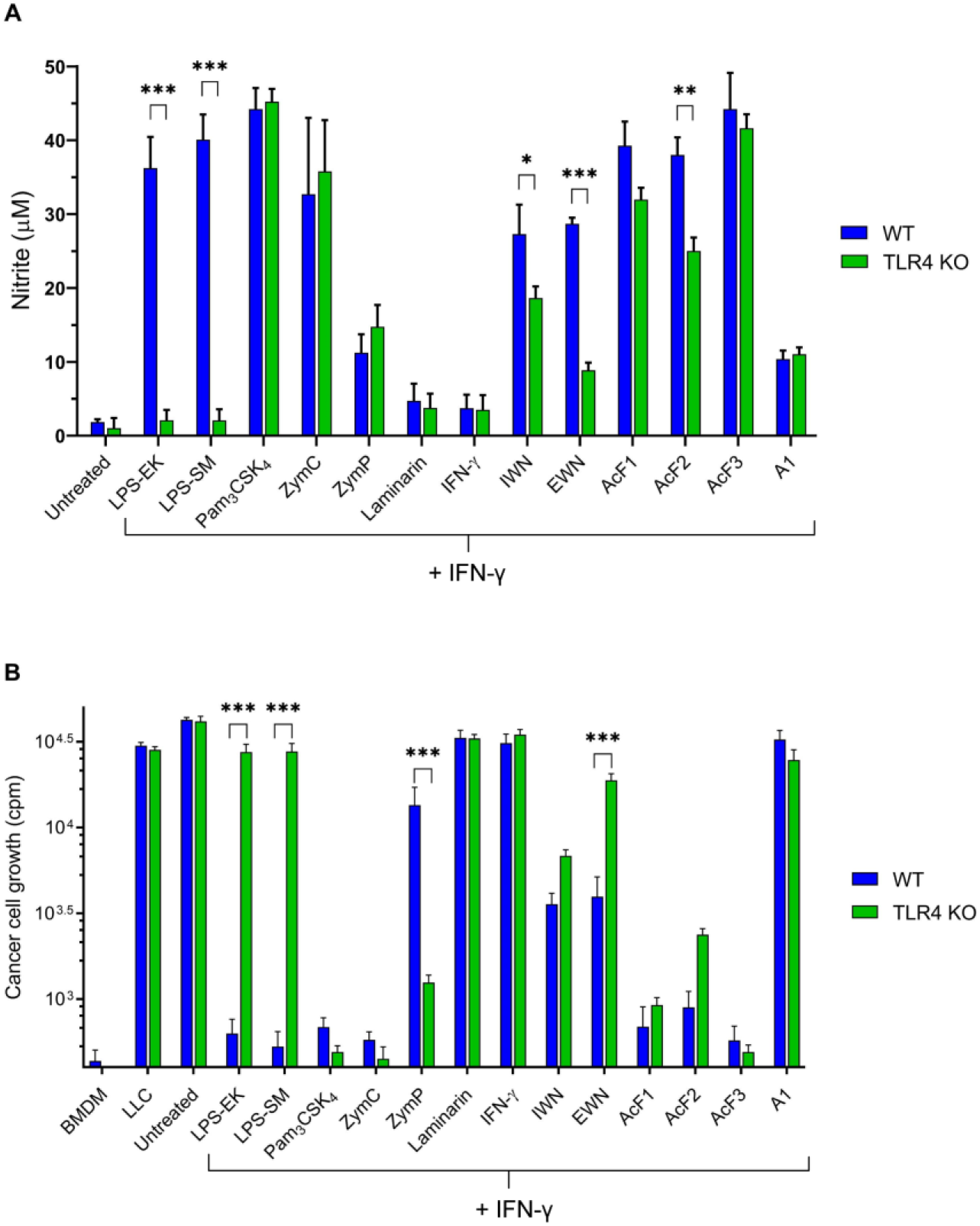
Polysaccharides from *I. obliquus* synergize with IFN-γ to produce NO in wild type (WT) and TLR4 knockout (KO) BMDMs, with subsequent growth inhibition of Lewis lung carcinoma (LLC) cells. BMDMs were incubated with mitomycin C for 2 h, before treatment with polysaccharides and IFN-γ (20 ng/mL) for 24 h. The *I. obliquus* polysaccharide concentration was 100 μg/mL. LPS-EK, LPS-SM, Pam_3_CSK_4_ (all 100 ng/mL), zymosan crude (ZymC), zymosan purified (ZymP) and laminarin (all 100 μg/mL) together with IFN-γ were used as controls. **A**) NO in the cell supernatants was quantified using the Griess assay, prior to the growth inhibition assay. **B**) Growth inhibition of LLC cells by activation of macrophages.. LLC cells (3 x 10^3^/well) were added to BMDMs (3 x 10^4^ cells/well) after activation. Only the results from the 10:1 ratio are shown. After 24h, radiolabelled thymidine was added and the cells were incubated for another 24h before inhibition of cancer cell growth was measured by radioactive counts per minute (cpm) for each sample. Three independent experiments were performed, and average values ± SD are shown. Statistical significance indicates the difference between WT and TLR4 KO cells, and was calculated using two-way ANOVA followed by Sidak’s multiple comparison test, *** = p < 0.001, ** = p < 0.01, * = p < 0.05.

### Several of the polysaccharides induce production of the pro-inflammatory cytokines IL-6 and TNF-α in human and mouse macrophages

In order to investigate if *I. obliquus* polysaccharides could also activate human macrophages, human monocytes-derived macrophages were stimulated with polysaccharides with and without IFN-γ for 24 h, before quantification of the pro-inflammatory cytokines IL-6 and TNF-α in the cell supernatants. Mouse BMDMs were also set up in the same type of experiment. As shown in Figure 7, several of the polysaccharides were able to induce IL-6 and TNF-α production in both human and mouse macrophages, with some differences between the two species. In the mouse macrophages, AcF1, AcF2 and AcF3 statistically significant induced IL-6 and TNF-α production, while IWN statistically significant induced IL-6 production but not TNF-α. Notably, AcF3 induced 5-10 fold more IL-6 and TNF-α than AcF1 and AcF2, comparable to the positive control Pam_3_CSK_4_. In the human macrophages, AcF3 was still the most active fraction for IL-6 production, but for TNF-α there were only small differences between the polysaccharides. The results demonstrate the pro-inflammatory potential of *I. obliquus* polysaccharides, with AcF3 standing out from the rest as being able to activate both human and mouse macrophages to produce high levels of IL-6 and TNF-α.

**Figure 7:**
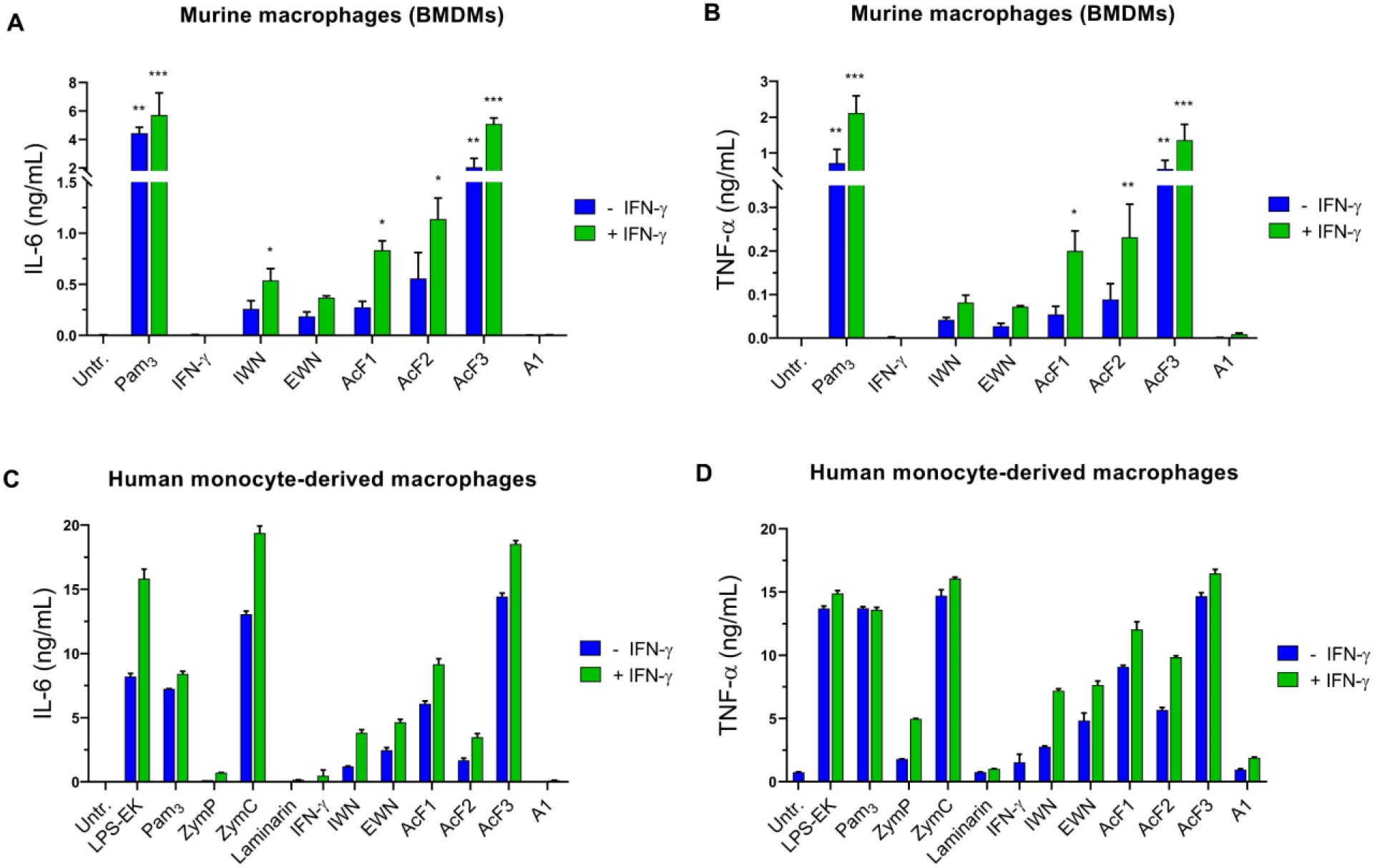
Polysaccharides from *I. obliquus* increased levels IL-6 and TNF-α production in human and mouse macrophages. Macrophages were treated with polysaccharides (100 μg/mL) alone or with IFN-γ (20 ng/mL) for 24h, before cytokine concentration in cell media was quantified using Luminex technology. **A** and **B**) Mouse BMDMs (2.5 x 10^5^ cells/well in 24-well plates) were treated with mitomycin C for 2h prior to activation with polysaccharides, before IL-6 (**A**) and TNF-α (**B**) were quantified in the cell supernatants. Pam_3_CSK_4_ (100 ng/mL) with or without IFN-γ was used as a positive control. Three independent experiments were performed, and average values ± SD are shown. Statistical significance was calculated using one-way ANOVA followed by Dunn’s comparison test, with single comparisons against the untreated control *** = p < 0.001, ** = p < 0.01, * = p < 0.05. **C** and **D**) Human monocyte-derived macrophages (1 x 10^5^ cells/well in 48-well plates) were treated with polysaccharides for 24h before IL-6 (**C**) and TNF-α (**D**) were quantified in the cell supernatants. LPS-EK, Pam_3_CSK_4_ (both 100 ng/mL), zymosan crude (ZymC), zymosan purified (ZymP) and laminarin (all 100 μg/mL), were used as controls. Two independent experiments were performed, and average values ± SD from one representative experiment are shown.

### The polysaccharides bind to various pattern-recognition receptors, including TLR2, TLR4 and Dectin-1a

To investigate which receptors could be responsible for the activation observed when treating macrophages with *I. obliquus* polysaccharides, HEK293 reporter cell lines transfected with specific PRRs were utilized. First, we wanted to investigate the interaction with TLR4 from the polysaccharide fractions that had reduced activity in the TLR4 KO cells. To test for hTLR4 interaction, samples were mixed with the LPS inhibitor PMB prior to cell stimulation. Quite astonishingly (due to their activity in the TLR4 KO macrophages), AcF1 and AcF3 were the most potent TLR4 agonists of the *I. obliquus* polysaccharides, and the activity was retained after treatment with PMB (Figure 8A). EWN and AcF2 were also able to activate TLR4, although to a lower degree, and the activity was gone at 10 μg/mL. Interestingly, whereas PMB-pre-treated LPS-EK in concentration 10 and 1 ng/mL showed no TLR4 activation, LPS-SM was only marginally inhibited at both concentrations. However, at 1 ng/mL the activity of non-pre-treated LPS-SM was very low compared to LPS-EK, suggesting that LPS-SM is a less potent TLR4 ligand while at the same time being less affected by PMB than LPS-EK.

**Figure 8:**
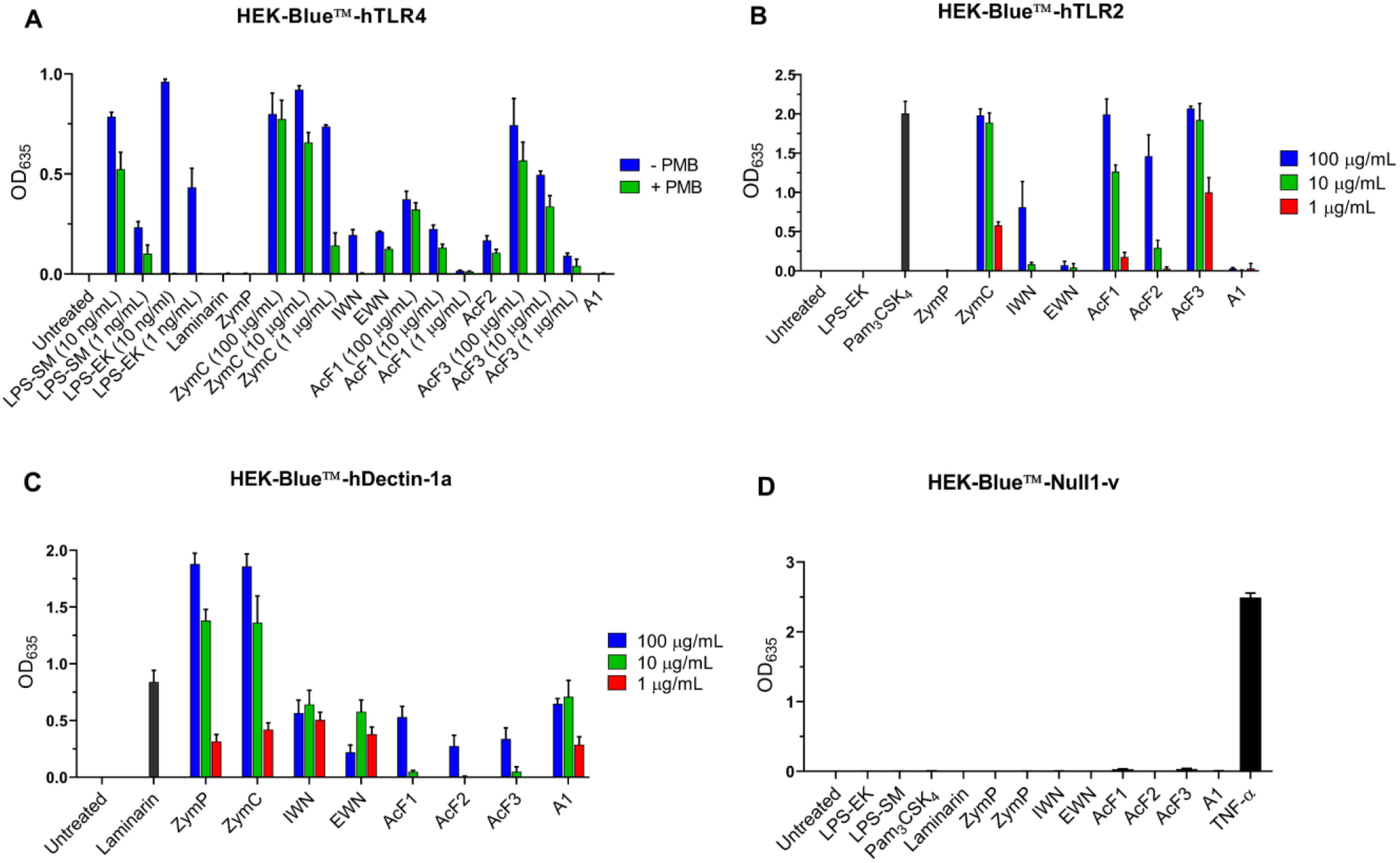
*I. obliquus* polysaccharides are agonists for the human immune receptors TLR4 (**A**), TLR2 (**B**) and Dectin-1a (**C**). HEK-Blue™ cells were incubated with polysaccharides or controls for 16 h. Then, OD635 was measured in order to detect secreted alkaline phosphatase (SEAP) in the supernatant. Polysaccharides were used at 1, 10 and 100 μg/mL; for the null cells, polysaccharides were used at concentration 100 μg/mL. LPS-SM, LPS-EK (10 and 1 ng/mL), Pam_3_CSK_4_ (100 ng/mL), laminarin (100 μg/mL), zymosan crude (ZymC) and zymosan purified (ZymP) (both 1, 10 and 100 μg/mL) and TNF-α (10 ng/mL) were used as controls depending on which reporter cell line was used. PMB concentration was 10 μg/mL for **A**. Three independent experiments were performed, and average values ± SD are shown.

Next, we wanted to investigate if the polysaccharides activated TLR2 (Figure 8B), since several publications suggest that fungal and plant polysaccharides are able to activate macrophages through this receptor (26–28). AcF1 and AcF3 stimulated hTLR2 strongly. EWN and A1 were completely inactive, while AcF2 and IWN were only active at the highest concentration tested (100 μg/mL). Interestingly, zymosan lost its TLR2-binding capability when used as the purified “β-glucan only” formulation, whereas the crude formulation containing mannan, proteins and other structures had potent activity that was comparable to AcF3 (29).

Because β-glucans of various kinds are known to bind to the C-type lectin receptor Dectin-1 (11, 30), we wanted to examine if the *I. obliquus* polysaccharides were agonists for this receptor as well. Specifically, a HEK-Blue™ reporter cell line expressing the Dectin-1a isoform was used, since this isoform can be activated by both soluble and particulate β-glucans in contrast to the HEK-Blue™ Dectin-1b reporter cell line is only activated by particulate β-glucans (31). As seen from Figure 8C, all six polysaccharides interacted with the human Dectin-1a (hDectin-1a) receptor, to various degrees. Interestingly, IWN, EWN and A1 were the most potent hDectin-1a ligands, whereas being the least active in the NO-, growth inhibition- and cytokine assays. The acidic polysaccharides AcF1, AcF2 and AcF3 gave detectable hDectin-1a activation only at the highest concentration tested (100 μg/mL). All the polysaccharides gave lower activation of hDectin-1a compared to the particulate β-glucan zymosan, but had comparable activity to the soluble β-glucan laminarin. None of the polysaccharides induced SEAP production in HEK293-null cells, indicating that the results are not due to activation of other receptors expressed by HEK293 cells (Figure 8D). Taken together, the results suggest that AcF1 and AcF3 are the most active TLR agonists of the isolated *I. obliquus* polysaccharides; they were able to bind to both TLR2 and TLR4 and to some extent Dectin-1a, but macrophage activation by these polysaccharides did not require activation of TLR4. The interaction with the receptors and the most important intracellular signalling pathways leading to production of pro-inflammatory mediators in the macrophages are summarized in Figure 9. In addition, Table 2 gives an overview of the immunomodulating activities of the *I. obliquus* polysaccharides used in this study.

**Figure 9:**
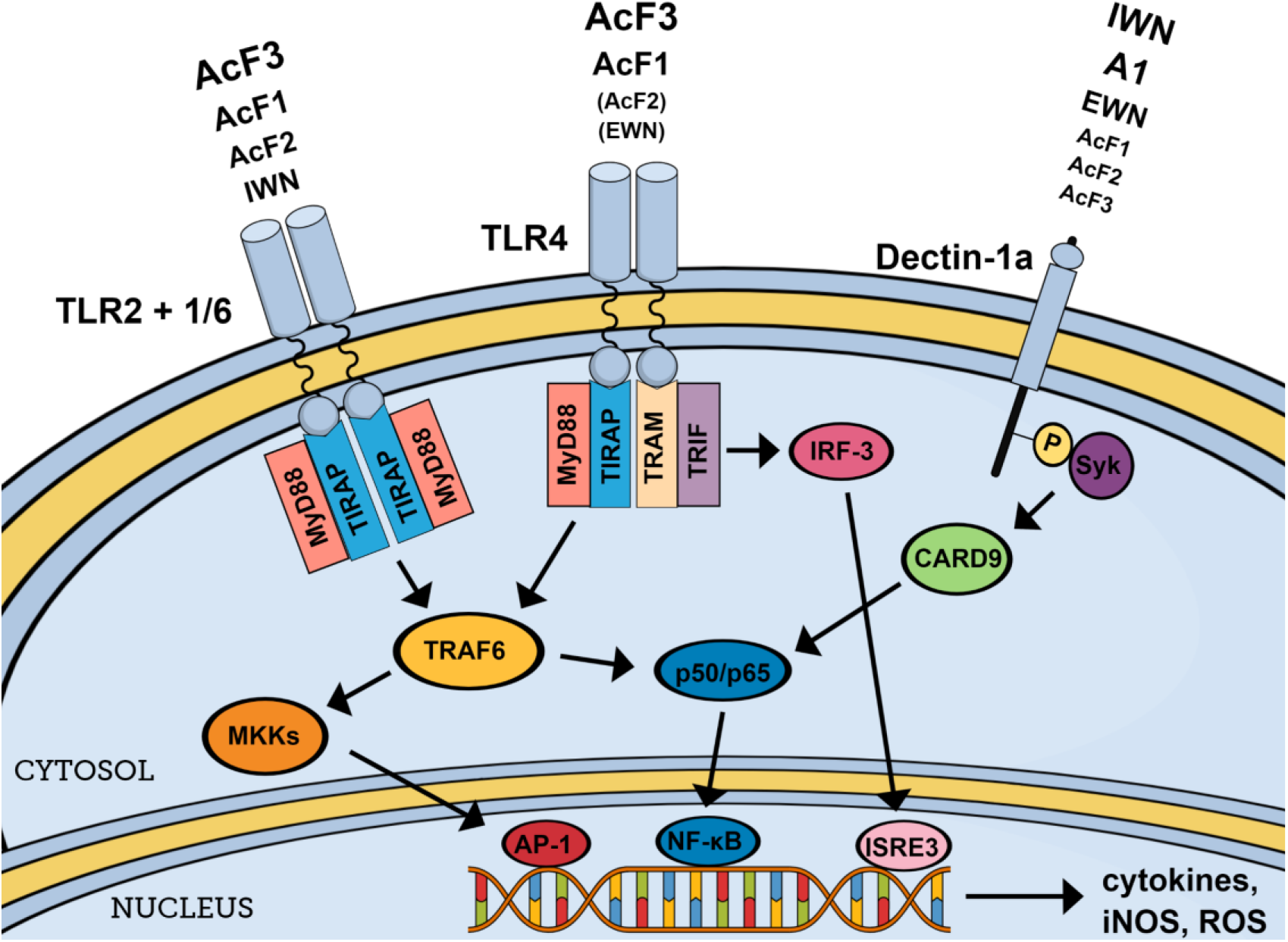
*I. obliquus* polysaccharides, in particular AcF1 and AcF3, are able to activate macrophages through TLR2 and TLR4. In addition, IWN, A1, EWN and to some extent AcF1, AcF2 and AcF3 activate Dectin-1a. The activation of these receptors could trigger production of pro-inflammatory cytokines and NO, which again may lead to functional outcomes such as growth inhibition of cancer cells.

**Table 2:**
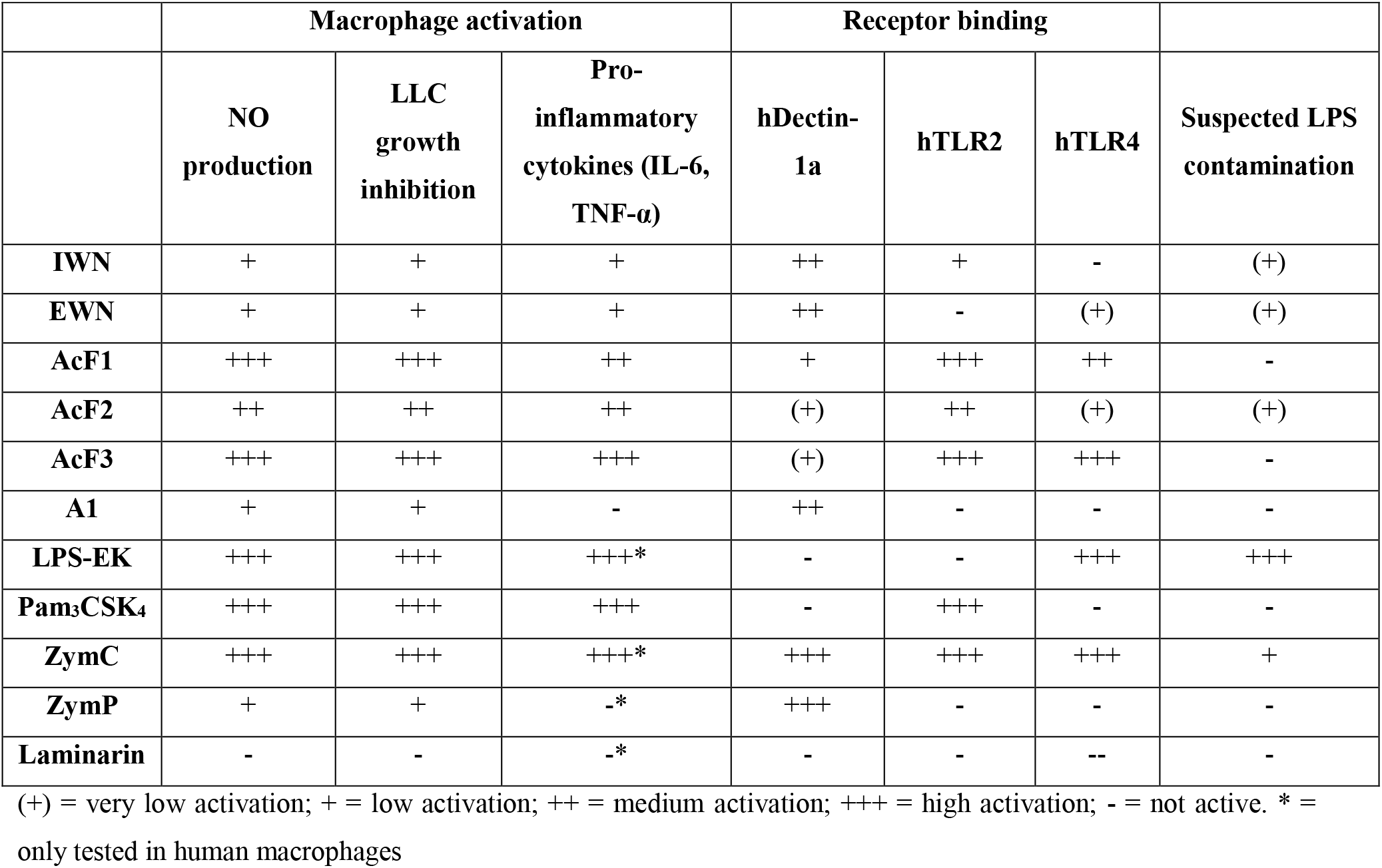
Summary of immunomodulating properties of *I. obliquus* polysaccharides and PRR ligands used in this study, and their interaction with the pattern recognition receptors dectin-1, TLR2 and TLR4. The macrophage activation is based on synergy with IFN-γ.

## Discussion

In this paper, we investigated and characterized immunological properties of six polysaccharides isolated from the medicinal fungus *I. obliquus*. Based on their structural motifs and molecular weight, the polysaccharides could be allocated to three main structural types (Table 1), defined as neutral and water-soluble (IWN and EWN), acidic and water-soluble (AcF1, AcF2 and AcF3) or neutral and particulate (A1). In the immunological assays used in this study, the acidic polysaccharides appeared to be more potent than the neutral polysaccharides. All polysaccharides were however able to induce nitric oxide production by mouse bone marrow-derived macrophages (BMDMs) by synergistic action with IFN-γ. In addition, all of the polysaccharides except A1 were able, in combination with IFN-γ, to induce growth inhibition of cancer cells through activation of the macrophages, which correlated with the levels of NO induced (Fig 2 and 5).

Furthermore, the acidic polysaccharides, in particular AcF3, induced potent production of IL-6 and TNF-α by human and mouse macrophages. The highly similar responses in the human and mouse cells suggest that the two cell types were activated through similar mechanisms and that the receptors responsible for activation are likely to be the same, which is also reflected by the high sequence similarities between TLRs from humans and mice (32). However, in preliminary experiments we were not able to induce nitric oxide (NO) production by the human macrophages after activation with the polysaccharides, in contrast to the mouse macrophages. It is well known that iNOS activation and subsequent NO production is difficult to achieve *in vitro* using human monocyte-derived macrophages (33). However, there seems to be little doubt about the importance of NO produced by macrophages in human physiology and immunology (34). Rather than being a reflection of the *in vivo* situation, the inability of human macrophages to produce NO *in vitro* could be explained by factors such as using sub-optimal sources of macrophages for this specific purpose or using the wrong differentiation/activation factors, leading to inactivation of the iNOS gene (20, 33). Because sufficient production of NO seems to be a requirement for growth inhibition of cancer cells by activated macrophages (8), the discrepancies in NO production between the human and mouse macrophages should be explored further.

The activity observed from the acidic, water-soluble polysaccharides correlated well with the ability to activate TLR2 and/or TLR4 in the HEK293 reporter cell lines. The main difference in terms of monosaccharide composition between the acidic and neutral polysaccharides was the presence of (1→4)-α-galacturonic acid in the acidic polysaccharides (12-17 % of total carbohydrate content). It therefore seems probable that this motif is important for the activation of macrophages and for the TLR2- and TLR4-binding capabilities. Interestingly, citrus pectin containing mainly (1→4)-α-galacturonic acid has previously been shown to bind both TLR2 and TLR4 (35). Another study found that citrus pectin containing de-esterified galacturonic acid was able to block the TLR2/1 heterodimer due to negative charges on the polymer associating with positively charged areas of TLR2, and that TLR2/6 activity was unaltered by the same polymer (36). These findings suggest that galacturonic acid is important for binding the receptor but is not necessarily responsible for activation of the receptor. Thus, it might be that other parts of the *I. obliquus* polysaccharides are important for TLR activation as well, requiring both galacturonic acid and some other structural motifs present in the acidic polysaccharides, such as β-glucose-containing motifs. However, the neutral polysaccharide IWN also activated TLR2 to some extent, whereas EWN activated TLR4 to some extent, although these fractions did not contain galacturonic acid. This suggests that the presence of galacturonic acid was not the only reason for the activation of TLR2 and TLR4. It is likely that some of the differences in activity are caused by variations in the three-dimensional structure of the polysaccharides, which is known to be important for polysaccharide interaction with various receptors (30). This could explain why AcF2 appeared less active at binding TLR2 and TLR4 than AcF1 and AcF3. According to GC-MS data, AcF2 exhibited a higher degree of side-chains than AcF1 and AcF3, which could affect the accessibility of the galacturonic acid. Efforts should be put into finding out which parts of the polysaccharides are responsible for the activity, for example by using enzymatic methods to degrade and isolate specific parts, and by investigating the three-dimensional structure of the polymers.

In addition to the synergetic activation with IFN-γ, AcF1 and AcF3 were able to induce some NO production when used alone, in contrast to the other polysaccharides, which required co-stimulation from IFN-γ. This might be explained by their ability to act through TLR4 as well as TLR2. Of the various extracellular TLRs, TLR4 is unique by it capability to induce iNOS in mouse macrophages without co-stimulation with other factors. It has been suggested that this characteristic comes from intracellular signalling through TRIF in addition to MyD88, leading to subsequent production of type I IFNs, which ultimately will function as an autocrine co-stimulatory signal on the macrophages (37).

To rule out that the activation through TLR4 was due to LPS contamination of the polysaccharide fractions, several experiments were conducted. First, LPS detection by a GC-MS assay concluded that the fractions contained less than 1.4 ng/mL LPS (19). Next, PMB treatment and subsequent macrophage activation demonstrated than activity from LPS-SM and LPS-EK at 1 ng/mL was completely blocked by PMB treatment, while at 10 ng/mL the activity was negligible. In contrast, the polysaccharide fractions were not affected by PMB treatment (Figure 5 and 8A). Thus, it was concluded that the activity of the polysaccharide fractions was not due to LPS contamination. In the TLR4 reporter cell line, LPS-EK and LPS-SM were included as controls at concentrations of 10 and 1 ng/mL. These were able to activate TLR4, and LPS-EK was completely inhibited by PMB at both concentrations. However, LPS-SM was only marginally inhibited (Figure 8A). This is a surprising finding as PMB is thought to inhibit and block functioning of LPS through the Lipid A part, which is present in both LPS-EK and LPS-SM. It might be that small variations in this region determine the efficiency of PMB, and this should be explored further, since most studies that use PMB as an LPS antagonist use LPS from *E. coli*. However, the results are not in agreement with the results from the macrophages, where PMB was able to block LPS-SM. It might be that the HEK-Blue cells are simply more sensitive to LPS due to for example higher receptor density, causing them to react to some types of LPS regardless of their association with PMB. Nevertheless, at a concentration of 1 ng/mL the activity of PMB-treated LPS-SM in the TLR4 reporter cell line was very low, and is thus unlikely to have caused the potent activation seen from AcF1 and AcF3.

Some authors have suggested that various plant and fungal extracts are contaminated with microbial lipoproteins that cause potent activation of TLR2, and that immunological activity observed in these studies stems from these lipoproteins rather than the isolated plant or fungal polymers (28, 38). To rule out the possibility of such a lipoprotein contamination in the *I. obliquus* extracts, we conducted an elemental combustion analysis to quantify the presence of nitrogen and carbon in our extracts, since bacterial lipoproteins contain large amounts of nitrogen readily detected by such elemental analysis. While one of our ethanol extracts contained significant amounts of nitrogen, none of the polysaccharide water extracts contained any traces of this element, strongly suggesting that our purified polysaccharides did not contain such a contamination (data not shown).

In addition to TLR2 and TLR4, some of the *I. obliquus* polysaccharides were able to bind to hDectin-1a. Dectin-1 is one of the most important and well-characterized β-glucan receptors on macrophages, recognizing (1→3/1→6) β-glucan with high affinity depending on polymer size, three-dimensional shape and degree of branching (39). Because all the *I. obliquus* polysaccharides had a main structural motif resembling fungal β-glucans, their ability to bind Dectin-1 was not surprising. However, the acidic polysaccharides showed low interaction with Dectin-1a compared to the immunologically less active fractions IWN, EWN and A1, which suggests that this receptor was not crucial for the immunological activity observed. Since Dectin-1 specifically recognizes β-glucan motifs, it is likely that the β-glucan parts of the polymer in IWN, EWN and A1 were more exposed to the outer environment compared to AcF1, AcF2 and AcF3, thus leading to higher affinity for Dectin-1a. The affinity toward Dectin-1a was lower for all isolated polysaccharides compared to the positive control zymosan, but when compared to laminarin, the activity was similar, suggesting that the differences in activity are due to different sizes and three-dimensional shapes rather than the β-glucan motif alone. There are many unknowns when it comes to the structure-activity relationship for Dectin-1 activation by β-glucans. It has been reported that particulate, large-sized β-glucans such as zymosan and lentinan are able to induce immune responses upon activation of Dectin-1, while water-soluble, smaller β-glucans like laminarin have antagonistic effects on the same receptor (13). Several isoforms of Dectin-1 exist in both humans and mice, with the two main types being a full-length Dectin-1a isoform that can recognize both soluble and particulate β-glucans, and a “stalkless” Dectin-1b isoform that recognizes only particulate β-glucans (31, 40). In addition, Dectin-1 has been reported to collaborate with both TLR2 (41) and TLR4 (42) on the macrophage cell membrane upon recognition of fungal structures, and the co-activation may give synergistic effects (43), further complicating the potential effects of complex polysaccharides such as the ones isolated from *I. obliquus* in this study. Thus, because all the isolated polysaccharides showed some affinity for Dectin-1a, its importance for the immunological activity cannot be ruled out, and further effort should be put into unravelling the relationship between the various receptors upon recognition of the *I. obliquus* polysaccharides. While Dectin-1 collaboration with TLRs is usually caused by the presence of multiple distinct effector molecules in crude formulations, such as mannoproteins, lipoproteins and β-glucans in crude zymosan (14), it could well be that the complexity of the *I. obliquus* polysaccharides leads to activation of several receptors using just “one molecule”. This could have interesting implications for the usage of such molecules as immunomodulating drugs. In addition, the high water-solubility of the polysaccharides might be attractive for *in vivo* situations compared to the particulate β-glucan formulations most often used in the literature. For example, in addition to having potential local effects in the gastrointestinal tract, smaller, water-soluble polysaccharides of fungal origin may be internalized by gut epithelial cells and systemically distributed shortly after ingestion as opposed to larger, particulate polysaccharides that require phagocytic uptake by macrophages (42).

In conclusion, we here report the immunological activity of several polysaccharides isolated from *I. obliquus*, with the acidic, water-soluble polysaccharides AcF1 and AcF3 in particular being able to induce a potent pro-inflammatory, tumoricidal phenotype in mouse macrophages. In addition, all polysaccharides except A1 were able to induce IL-6 and TNF-α production by human macrophages. The activation was likely induced through binding of TLR2, TLR4 and Dectin-1a, and the tumoricidal potential of the polysaccharides through interaction with multiple receptors might provide interesting new opportunities for cancer immunotherapy.

